# Unleashing meiotic crossovers in hybrid plants

**DOI:** 10.1101/159640

**Authors:** Joiselle Blanche Fernandes, Mathilde Seguéla-Arnaud, Cecile Larchevêque, Andrew H. Lloyd, Raphael Mercier

**Affiliations:** Institut Jean-Pierre Bourgin, INRA, AgroParisTech, CNRS, Université Paris-Saclay, RD10, 78000 Versailles, France

## Abstract

Meiotic crossovers shuffle parental genetic information, providing novel combinations of alleles on which natural or artificial selection can act. However, crossover events are relatively rare, typically one to three exchange points per chromosome pair. Recent work has identified three pathways limiting meiotic crossovers in *Arabidopsis thaliana*, that rely on the activity of FANCM^1^, RECQ4^2^ and FIGL1^3^, respectively. Here, we analyzed recombination in plants where one, two or three of these pathways were disrupted, in both pure line and hybrid contexts. The highest effect was observed when combining *recq4* and *figl1* mutations, which increased the hybrid genetic map length from 389 to 3037 centiMorgans. This corresponds to an unprecedented 7.8-fold increase in crossover frequency. Disrupting the three pathways do not further increases recombination, suggesting that some upper limit has been reached. The increase in crossovers is not uniform along chromosomes and rises from centromere to telomere. Finally, while in wild type recombination is much higher in male than in female meiosis (490 cM vs 290 cM), female recombination is higher than male in *recq4 figl1* (3200 cM vs 2720 cM), suggesting that the factors that make wild-type female meiosis less recombinogenic than male wild-type meiosis do not apply in the mutant context. The massive increase of recombination observed in *recq4 figl1* hybrids opens the possibility to manipulate recombination to enhance plant breeding efficiency.

---

Crossovers (COs) are reciprocal exchanges between homologous chromosomes, with two consequences: First, in combination with sister chromatid cohesion, COs provide a physical link between homologs, which is required for balanced segregation of chromosomes at meiosis. Failure in the formation of at least one crossover per chromosome pair is associated with reduced fertility and aneuploidy^4^ Second, COs lead to the exchange of flanking DNA, generating novel mosaics of the homologous chromatids. This translates into genetic recombination, classically measured in Morgans (or centiMorgans cM). The number of COs appears to be constrained by both an upper and lower limit (Figure 1)^5^. As an illustration, Figure 1 plots the genetic size (cM) versus the physical size (DNA base pairs, log scale) of chromosomes in a diverse panel of eukaryotes. Note the large variations in physical size, while the genetic map length is much less variable: At the lower limit, chromosomes measure 50 cM, which corresponds to the requirement of at least one CO per chromosome pair (0.5 per chromatid). Across all species, most chromosomes do not receive many more than one CO per meiosis, 80% of them having less than three. High level of COs per chromosome, observed in handful of species such as S. *pombe*, appears thus to be counter-selected in most species, suggesting that CO rate is a trait under selection in both directions^6^.

**Figure 1.**
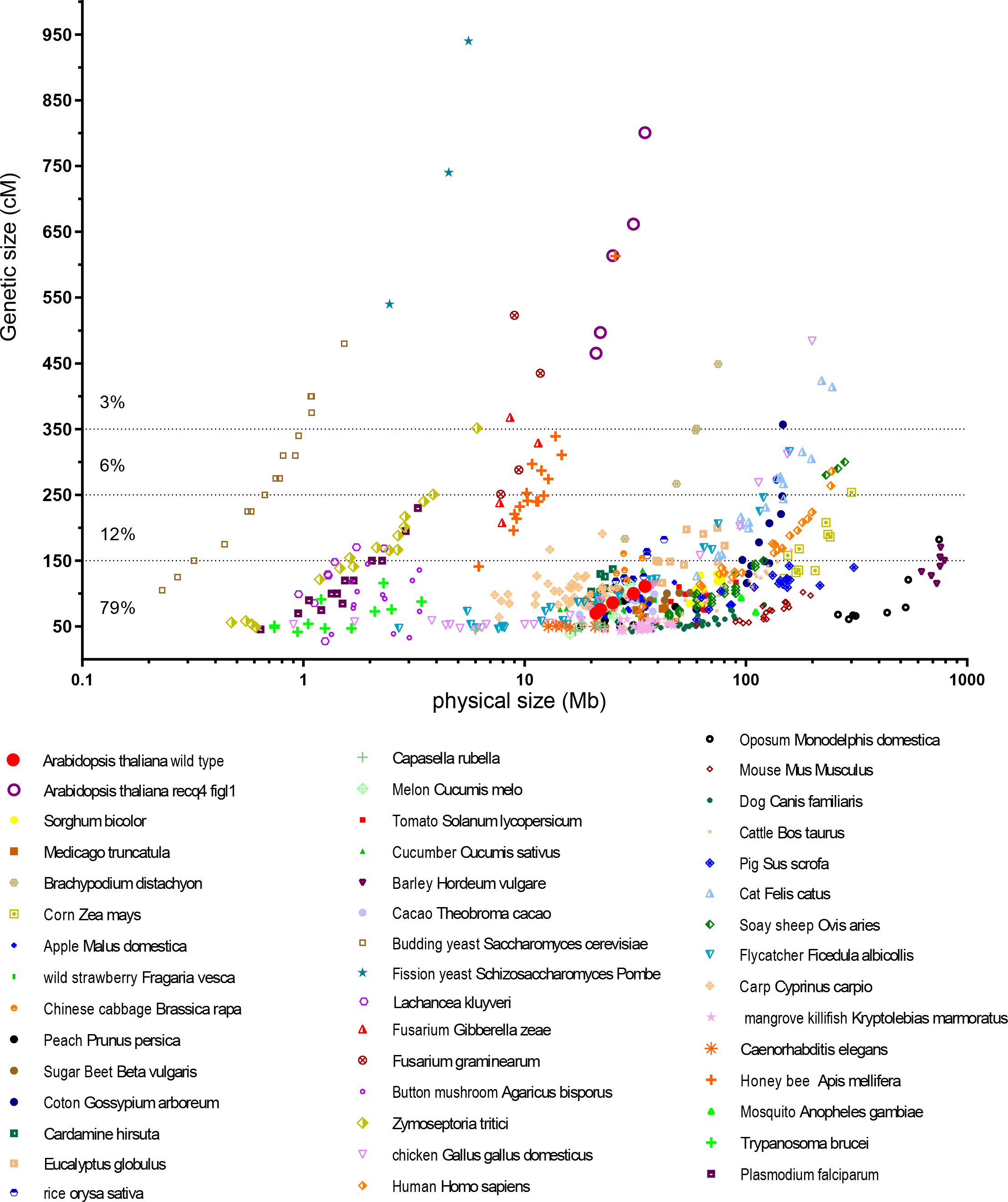
The number of meiotic crossovers is constrained in eukaryotes. Chromosomes from a range of species are plotted according to their physical (X axis, Mb, log scale) and genetic size (Y axis, cM, linear scale). Physical size is based on genome sequence assembly and genetic size is based on F2 or male/female average. Sex chromosomes have been excluded. An earlier version of this figure, with fewer species represented, was published in Mercier et al^5^.

Plant breeding programs rely on meiotic COs that allow the stacking of desired traits into elite lines. However, as described above, the number of COs is low. Further, some regions are virtually devoid of COs, such as the regions flanking centromeres^7^. This limits the genetic diversity that can be created in breeding programs and in addition, limits the power of genetic mapping in pre-breeding research. Increasing recombination is thus a desired trait in plant breeding^8,9^. Forward screen approaches have identified thee pathways that limit recombination in *Arabidopsis thaliana*, relying respectively on the activity of (i) FANCM helicase and its cofactors^1,10^, (ii) the BLM/Sgs1 helicase homologues RECQ4A and RECQ4B and the associated proteins TOP3α and RMI1^2,11^;(iii) the FIGL1 AAA-ATPase^3^. *RECQ4A* and *RECQ4B* are duplicated genes that redundantly limit meiotic crossovers in *Arabidopsis*, and will be herein designated as RECQ4, for simplicity. In all mutants, recombination is largely increased compared to wild type, when tested in pure lines. However, the *fancm* mutation, which increases meiotic recombination three fold in pure lines, has almost no effect on recombination in hybrids^3,12^. It thus remained unclear if the manipulation of these three pathways could be used to increase recombination in hybrids, the context that matters for plant breeding programs.

## Results

Here we analyzed recombination in single, double and triple mutants for *FANCM, RECQ4* and *FIGL1*, in both pure line and hybrid contexts, using two complementary approaches. First, we used a Fluorescent Tagged Line (FTL I2ab) that relies on transgenic makers conferring fluorescence to pollen grains organized in tetrads^13^ (Figure 2). With this tool we accurately measured recombination in two adjacent intervals in either pure line (Col) or F1 hybrid contexts (Col/Ler). Second, we used the segregation in F2 progeny of a set of 96 Col/Ler SNPs^3^ to measure recombination genome-wide in F1 hybrid contexts (Figure 3). In pure Col, each single mutant has higher recombination than wild type in both FTL tested intervals (Figure 2A-B). Each double-mutant is higher than the respective single mutants, confirming that *FANCM, RECQ4* and *FIGL1* act in three different pathways to limit COs. The highest increase is observed in the *fancm recq4* and *figl1 recq4* double mutants both with a ~10-fold increase compared to wild type. The observation that each double mutant combination results in an increase in recombination compared to the single mutants (i.e. three independent pathways), predicts that the triple mutant should even further increase recombination. However, recombination in the *figl1 recq4 fancm* triple mutant is not higher than in the highest doubles, suggesting that some upper limit has been reached. In the Col/Ler hybrid context (Figure 2C-D for FTLs; Figure 3A for genome wide genetic map), both *figl1* and *recq4* increased recombination, but *fancm* had no detectable effect as previously described in several hybrid contexts^3,12^. Similar to observations in pure Col, double-mutant hybrids have higher recombination than the corresponding singles. Notably, the *fancm* mutation leads to a detectable increase in recombination when combined with *figl1* or *recq4* (Figure 2C-D, Figure 3A). This predicts again that combining the three mutations should lead to a further increase. However, recombination is not statistically higher in the *figl1 recq4 fancm* triple-mutant compared to the *figl1 recq4* double and appears to be even reduced (Figure 2C and 3A).

**Figure 2.**
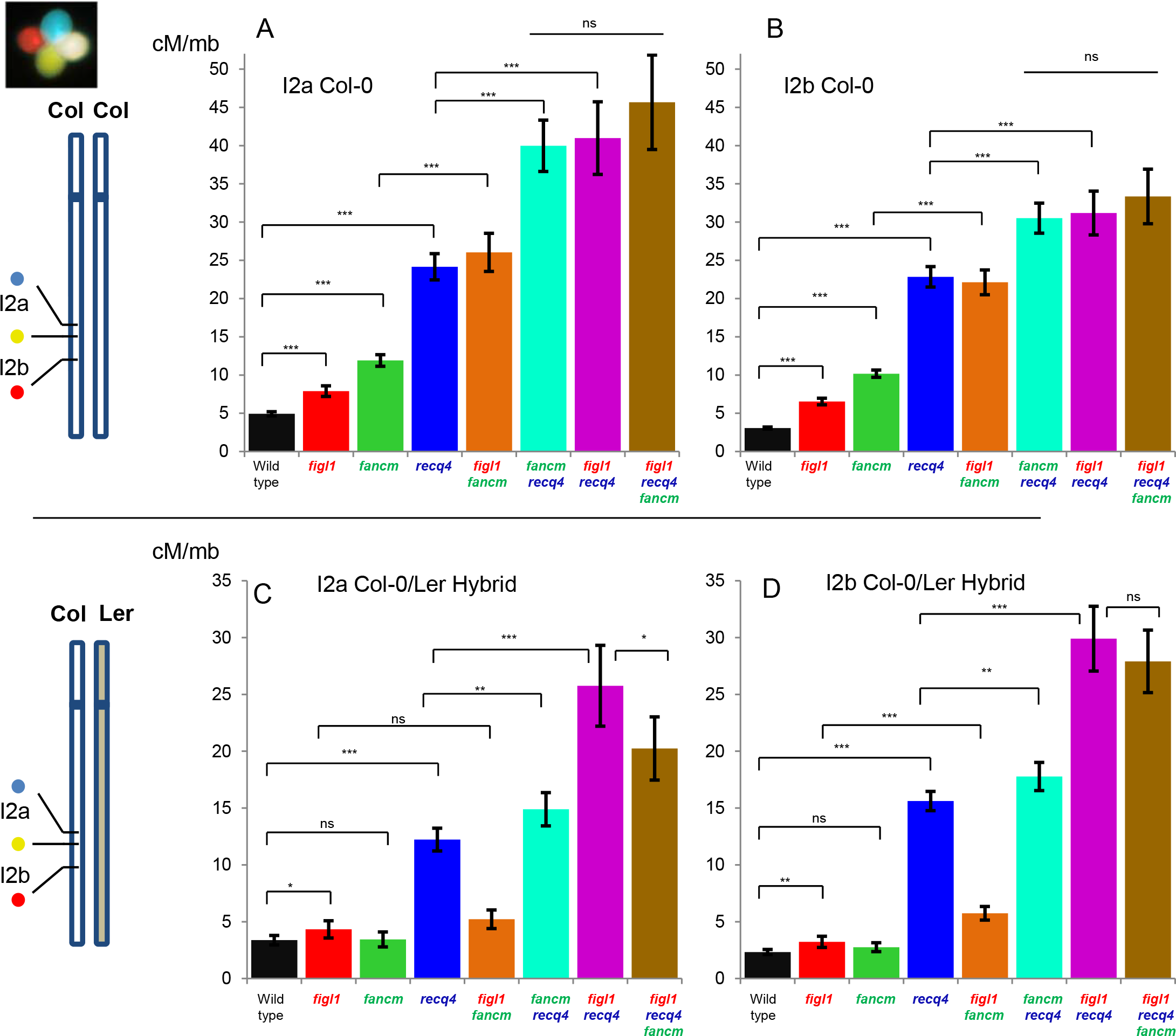
Meiotic recombination measured by FTL tetrad analysis. (A, B) recombination in a pure Col-0 line, in intervals I2a and I2b, respectively. (C, D) Recombination in the Col/Ler F1 hybrids, in intervals I2a and I2b, respectively. Bars are genetic distance calculated with the Perkins equation in cM/Mb, +/- 95% confidence interval. Z-tests are indicated: ***:p<0,001; **:p<0,01; *p<0.05; ns: p>0,05 Raw data can be found in table S1.

**Figure 3.**
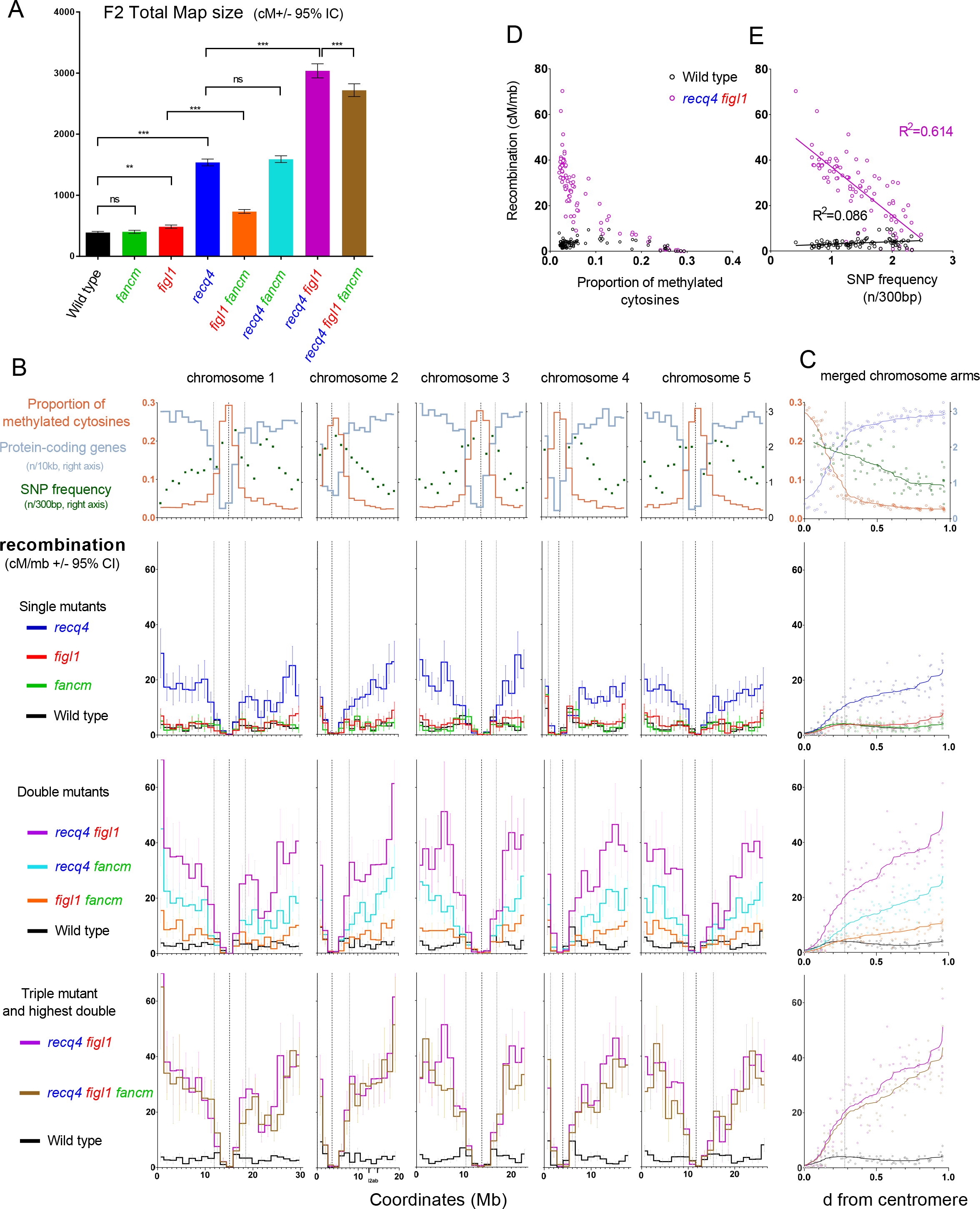
Genome wide recombination measured by SNPs segregation in F2s. (A) Total genetic maps. Tests are one-way-anova on the observed number of COs per F2 plant, with Dunnet correction. (B) Plotted along each of chromosomes (Mb), (from top to bottom) is gene density, single nucleotide polymorphism (SNP) density^28^, proportion of methylated cytosines and recombination in the different genotypes (cM/Mb, Haldane equation). Vertical dotted lines mark the position of the centromeres and peri-centromeres. (C) The same data merged for all chromosome arms (short arms of chromosome 2 and 4 have been excluded), plotted along relative distance form centromere, and smoothed. (D) Each interval is plotted according to the proportion of methylated cytosines and recombination frequency. (E) Each interval is plotted according to SNP density and recombination frequency. Linear regressions and associated R squares are shown.

The highest increase in the hybrid context is thus observed in the *figl1 recq4* double mutant, with a ~11-fold increase in the FTL I2ab intervals (Figure 2 C-D) and 7.8-fold increase genome wide (Figure 3A), from 389 ± 18 cM in wild type to 3037 ± 115 cM in *figl1 recq4* (which translates into 7.8 ± 0.4 and 60.7 ± 2.3 COs per meiosis, respectively). This is much higher than the highest increase in meiotic recombination observed so far in any mutant^14–16^. While *Arabidopsis* wild type has a very typical frequency of CO, with the genetic maps of each of the five chromosomes ranging from 70 to 110 cM (on average 1.6 CO per chromosome), the *figl1 recq4* is almost the highest recombining eukaryote (Figure 1) with the genetic maps of chromosomes ranging from 450 to 800 cM (on average 12 COs per chromosome: Figure 1).

We addressed if such a large increase in recombination could reduce the fertility of plants, precluding use of these mutant combinations in breeding programs. In hybrids, the number of seeds per fruit was not significantly reduced in any genotype (Figure 4B). However, a slight defect in pollen viability was observed in single and multi-mutants (Figure 4D). Further, in pure Col, both reduced seed set and pollen viability defects were detected, and were highest in *recq4 figl1* and *recq4 figl1 fancm* (Figure 4A, 4C). This suggests that some fertility defects may be associated with increased recombination. However, the fertility defects in the different genotypes are quite poorly correlated with the increases in COs (Compare Figure 2 and 4). For example, Col *fancm recq4* has the same large increase in recombination observed in *figl1 recq4* (Figure 2A-B), but pollen viability is much less affected (Figure 4C). This suggests that high levels of COs are not responsible *per se* for reduced fertility. We further examined meiosis in the Col *recq4 figl1* and *recq4 figl1 fancm* which show >40% pollen death (Figure 5). At diakinesis, homologous chromosomes are associated in pairs, connected by COs. The chromosomes appear more tightly connected in the mutants than in wild type, presumably because of increased CO numbers (Figure 5A-B). The shape of chromosomes at metaphase I, is also suggestive of increased CO numbers (Figure 5 C-D). At metaphase II, chromosome spreads revealed the presence of a few chromosome fragments and chromosome bridges in 50% of the cells (n=115 metaphase II cells; Figure 5 E-G), suggesting defective repair of a small subset of recombination intermediates. We propose that the absence of *FIGL1* and *RECQ4* disturbs the DSB repair machinery, leading to the formation of aberrant intermediates (such as the multi-chromatid joint molecules observed in the yeast *recq4* homologue *sgs1*^17^), most of them being repaired as extra-COs and a few failing to be repaired. We suggest that those unrepaired intermediates are responsible for most of the reduced fertility, but we cannot fully exclude that the extra COs themselves slightly disturb chromosome segregation.

**Figure 4.**
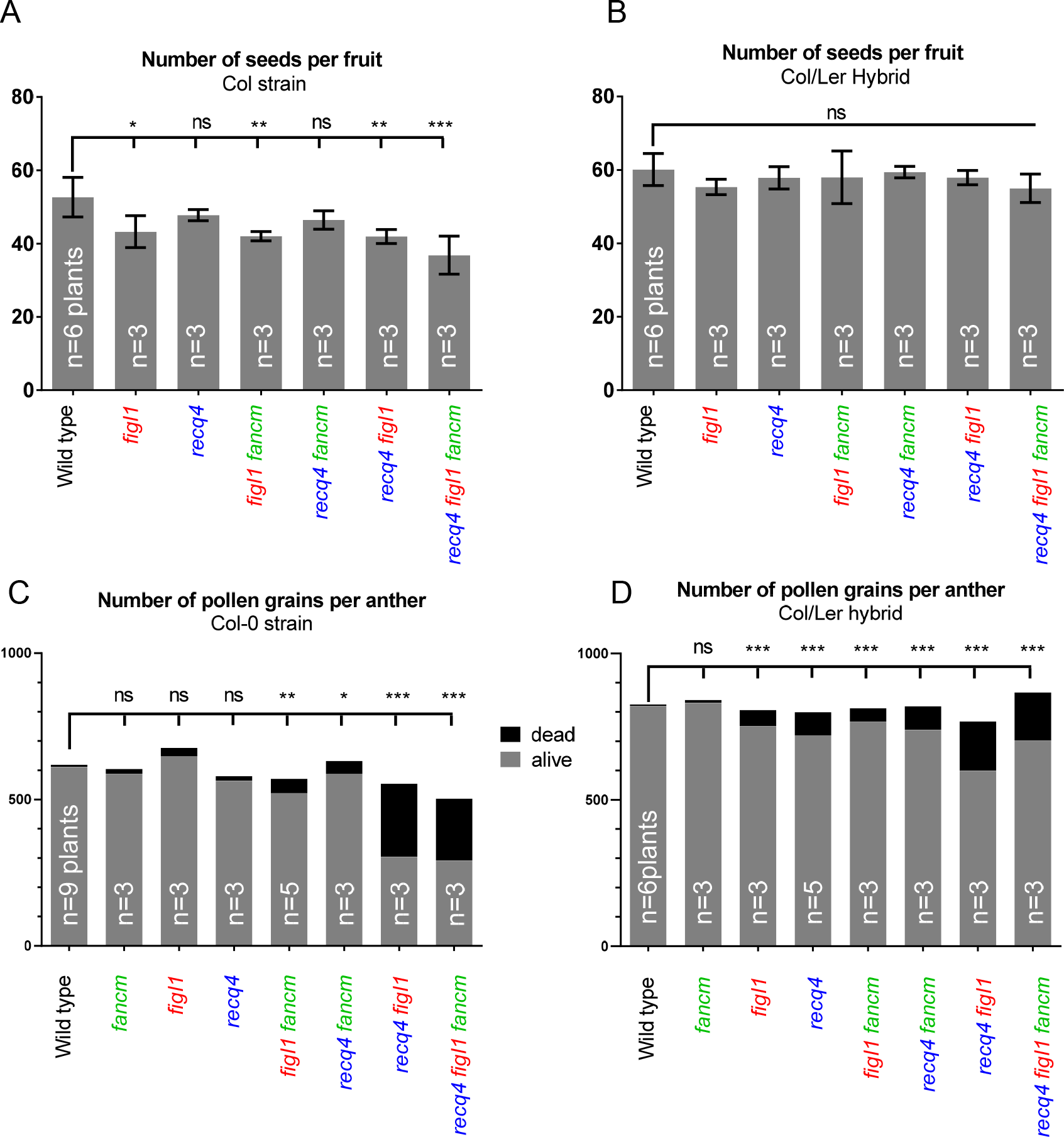
Analysis of fertility. (A, B) Number of seeds per fruit in pure Col and Col/Ler F1 hybrids, respectively. (C, D) Average number of viable and non-viable pollen grains per anther, following alexander staining. n indicates for the number of plants analyzed. For each plant, the number of seeds has been counted in a minimum of 10 fruits, and the number of pollen grains in a minimum of three anthers. Errors bars: +/- SD. Tests compare each genotype with the wild type, and are done by Anova followed by Dunnet correction for multiple tests.

**Figure 5.**
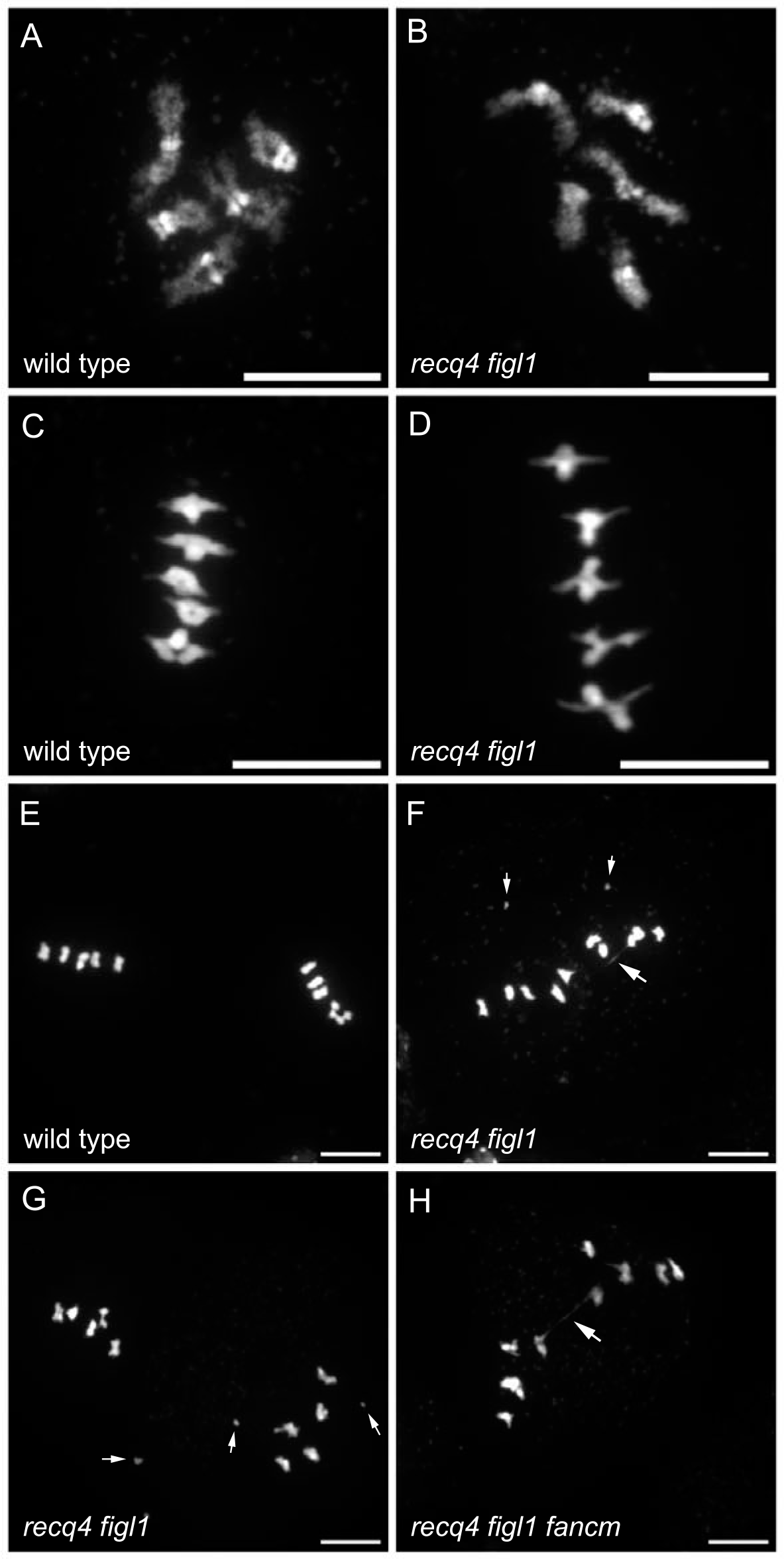
Meiotic chromosome spreads. (A, B) Diakinesis. Five chromosome pairs are observed in both wild type (Col) and *recq4 figl1* and appear to be more tightly linked in *recq4 figl1*. (C,D) Metaphase I. The five chromosome pairs are aligned on the metaphase plate. (E-H) Metaphase II. In wild type, five pairs of chromatids align on two metaphase plates. In *recq4 figl1* and *recq4 figl1fancm*, chromatid bridges (large arrows) and chromosome fragments (small arrows) are observed in 50% of the cells. Scale bars=10μM.

Next, we explored the distribution of COs along the genome (Figure 3 B-C) using the Col/Ler genome-wide recombination data. The first line of figure 3B-C shows gene density, single nucleotide polymorphism density^28^ and proportion of methylated cytosines^30^ plotted along each of chromosomes. The centromeres (vertical dotted line in figure 3B) are flanked by peri-centromeric regions having low gene density and high methylated cytosines (>7.5%, the genome average. Delimited by thin dotted lines). The three bottom lines of figure 3B-C show recombination along the chromosomes in single, double and triple mutants, respectively. At the scale used in this study (bins of ~1.3 Mb), the wild type curve of recombination frequency oscillates around ~2.5-3.5 cM/Mb in chromosome arms, with the maximums observed close to peri-centromeric and telomeric regions. The interval that spans the centromere in each chromosome, have very low levels of recombination. In accordance with results for total map size (figure 3A), *fancm* is indistinguishable from wild type. The curves of all the other mutant genotypes show higher recombination than wild type with *recq4 figl1* having the strongest effect (Figure 3 B-C). In all mutants, the same tendency is observed, with no effect on recombination for intervals encompassing or immediately flanking the centromere and an increase of the recombination with distance from the centromere to reach a maximum close to telomeres. In *figl1 recq4*, the recombination frequency rises rapidly from the centromere to the first third of the arm, where it reaches ~20 cM/Mb (~5-fold higher than wild type levels). This first third of the chromosome arms correspond to the peri-centromeric region, with a progressive decrease in methylation and increase in gene density with distance from centromere (Figure 3B-C, first row). In the remaining two third of the chromosomes, methylation and gene density is relatively stable, but recombination continues to increase towards the telomere, reaching an average of 45 cM/Mb (›10-fold higher than wild type levels). Knocking down *recq4* and *figl1* therefore has very different effects in different regions of the chromosome. We propose three non-exclusive possible causes for this effect. First, the position along the chromosome itself could influence CO frequencies, as recombination occurs in the context of highly organized and dynamic chromosomes^18^. Second, the accessibility of the chromatin may directly account for CO frequency, notably by influencing SPO11-dependant double strand break formation^19^. The anti-correlation of cytosine methylation and recombination (Figure 3D) seen in *recq4 figl1*, supports this view. Third, DNA polymorphism, which decreases from centromere to telomere (Figure 3C) may also prevent CO frequency. We observed a strong anti-correlation between recombination and SNP density in *recq4 figl1*, which does not exist in wild type (Figure 3E). As distance from centromere, methylation and polymorphisms are correlated with each other, it is difficult to decipher the relative importance of these three parameters for recombination in *recq4 figl1*. Interestingly, a drop in recombination is observed in the middle of the right arm of chromosome 1 in *recq4 figl1* and *recq4 figl1 fancm* (figure 3B), corresponding to a region of high polymorphism associated with a cluster of NBS-LRR disease resistance genes^20^. This supports the hypothesis that DNA polymorphisms discourage extra COs in the mutants. In some crops such as wheat^21^, barley^22^ or tomato^23^, large portions of chromosomes surrounding centromeres receive virtually no COs, albeit containing genes that are thus out of reach for breeding. It would be of particular interest to test the effect of *recq4 figl1* in such crops to see how the CO increase we see in the short peri-centromeres of Arabidopsis translates to their large peri-centromeres.

Next, we addressed whether the increase in recombination that we observed in F2s, equally affects male and female meiosis, by backcrossing the F1 hybrids (wild type, *recq4* or *recq4 figl1*) by Col-0, either as male or as female and analyzing SNP segregation in the progeny (Figure 6). In wild type the male genetic map was ~70% larger than in female (495±38 cM vs 295±28 cM, p<10^−4^), the difference being particularly marked in the vicinity of telomeres where male recombination is at its highest and female at its lowest (Figure 6B-C), in accordance with previous data^24^. In sharp contrast, in *recq4* and *recq4 figl1* the female maps are larger than the male maps (p<10^−4^; Figure 6A) and the CO distributions becomes similar in both sexes (Figure 6B-C). The total female map is increased from 295 ± 28 cM in wild type to 3176 ± 190 cM in *recq4 figl1*, a more than 10-fold increase. The most spectacular increase being observed at the vicinity of telomeres where the recombination rate is on average 0.8 cM/Mb in female wild type and 47 cM/Mb in female *recq4 figl1*. In wild type, the vast majority of COs are of class I while *recq4* and *figl1* mutations increase class II COs^2,3,5^. Accordingly, the effects of CO interference, which limits the occurrence of two adjacent class I COs, are observed genetically in wild type but not in the mutants with increased recombination (Table S1 and Figure 6D). It appears, therefore, that regulation of class I COs shapes their distribution along chromosomes and differentiates male and female meiosis, but this regulation does not apply to the class II COs up-regulated in the *recq4 figl1* double mutant.

**Figure 6.**
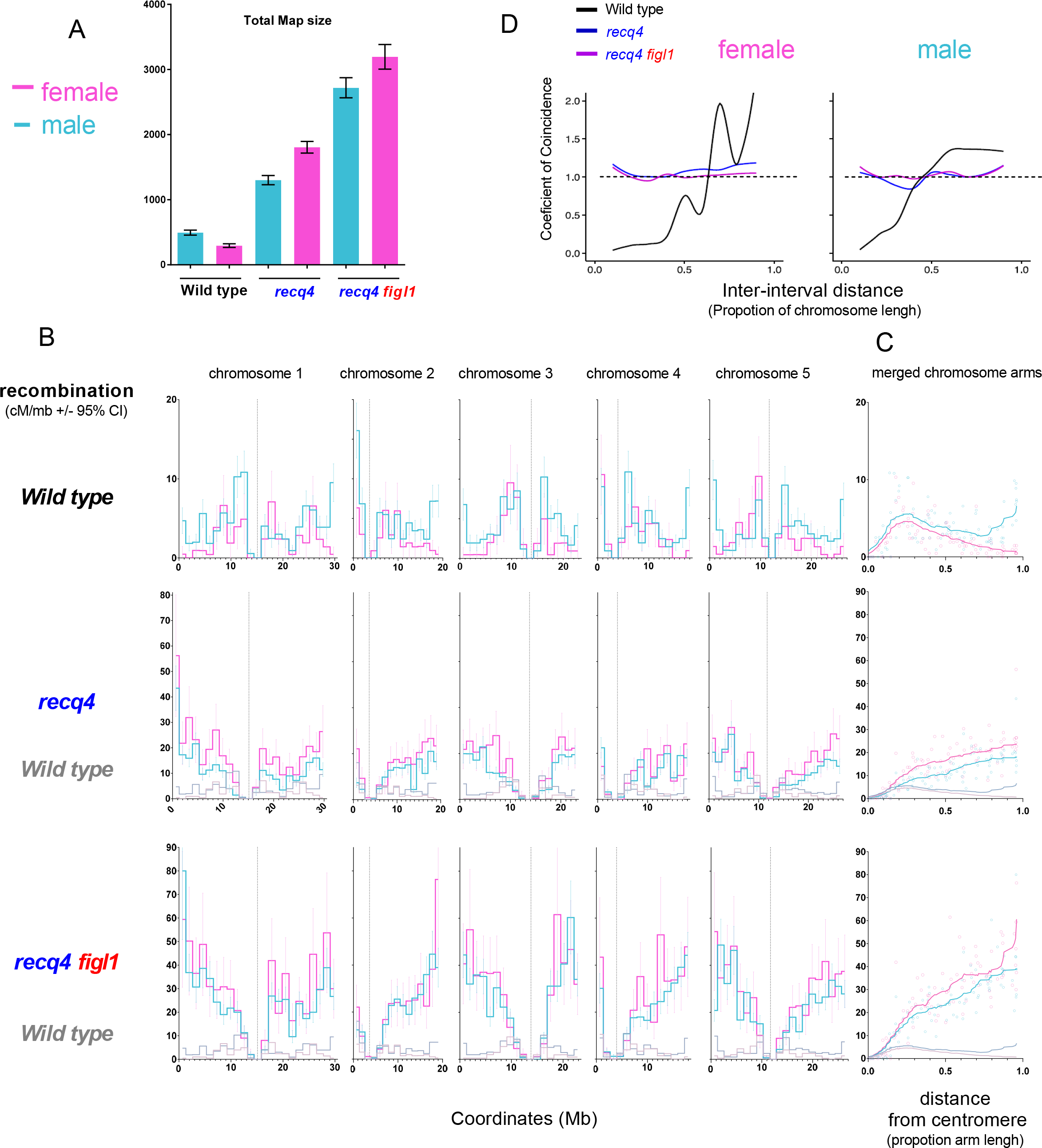
Male and female genome wide recombination. (A) Total male (blue) and female (pink) genetic maps in the wild type, *recq4* and *recq4 figl1*. (B) Plotted along each of the five chromosomes (Mb), is male and female recombination in the different genotypes (cM/Mb, Haldane equation). Errors bars represent 95% confidence intervals. Vertical dotted lines mark the position of the centromeres. (C) The same data as in B, merged for all chromosome arms (short arms of chromosome 2 and 4 have been excluded), plotted along relative distance form centromere and smoothed. (D) CoC curves. Coefficient of Coincidence is the ratio of the frequency of observed double COs to the frequency of expected double COs (the product of the CO frequencies in each of the two intervals). A CoC curve^31^ plots this ratio for each pair of intervals, as a function of inter-interval distance. In the presence of CO interference, CoC is around 0 for small inter-intervals distances, increasing to 1 with fluctuation around that value for larger inter-interval distances, as seen here for wild type. A flat line at 1, as observed for the mutants, suggests an absence of interference.

We showed that a large CO frequency is obtained when combining *recq4* and *figl1* mutations, resulting in a 7.8-fold increase in total map size, with limited effect on plant fertility. This opens the possibility to manipulate recombination in plant breeding programs, to increase shuffling of genetic information, break undesirable linkage, combine desirable traits, or increase the power of genetic mapping in pre-breeding research. It is intriguing that very few eukaryotes have high CO levels per chromosome while it is possible, as shown here and by a few exceptional eukaryotes. One possibility is that a high level of COs is associated with segregation problems at meiosis and thus fertility defects. Our data could support this possibility. Alternatively, it is possible that selection for a low level of recombination limits COs in eukaryotes, which could be optimal for adaptation in most contexts. An attractive idea is that in a stable environment, high recombination levels would break favorable allelic combinations that have been selected in previous generations^25^. The ZMM pathway that accounts for most COs in plants, mammals and fungi may have thus arisen in early eukaryotes to address the opposing constrains of ensuring at least one CO per chromosome while being able to adjust their numbers to low levels.

## Methods

The mutations used in this study were: Col: *fancm-1*^1^, *figl1-1*^3^, *recq4a-4* (N419423)^26^, *recq4b-2* (N511130)^26^; Ler: *fancm-10*^3^, *figl1-12*^3^, *recq4a-W387*^2^. The tetrad analysis line was I2ab (FTL1506/FTL1524/FTL965/*qrt1-2*)^13^. Hybrid lines were obtained through the crossing of Col plants bearing the *fancm, recq4a, recq4b, figl1, qrt* mutations and the FTL transgenes I2ab to Ler plants bearing the *fancm, recq4a, figl1and qrt* mutations. F1 plants were grown in growth chambers (16h/day 21°C, 8h/night 18°C, 65% humidity) and genotyped twice for each mutation (Table S5). F1 sibling plants of the desired genotypes were used for tetrad analysis, fertility measures, selfed to produce the F2 population and crossed as male or female to Col-0 to produce BC1 populations. Tetrad analysis (Figure 2), including data collection, measures of recombination (Perkins equation) and interference (Interference ratio) and statistical tests were performed as described in Girard *et al*^3^. F2 and BC1 populations were grown for three weeks and 100-150 mg leaf material was collected from rosettes. DNA extraction and genotyping for Col/Ler polymorphisms was performed using the KASPAR technology at Plateforme Gentyane, INRA Clermont-Ferrand, France. The set of 96 KASPAR markers (Table S4), which are uniformly distributed on the physical map (~every 1,3Mb) has been described in Girard *et al*^3^. Genotyping data were analyzed with the Fluidigm software (http://www.fluidigm.com) with manual corrections. The raw genotyping data set is shown in table S3. Recombination data were analysed with MapDisto 1.7.7.0.1.1^27^: Genotyping errors were filtered using the iterative error removal function (iterations=6, start threshold=0,001, Increase=0,001). Recombination (cM+/- SEM) was calculated using Classical fraction estimate and the Haldane mapping function. The Haldane function has been preferred to the Kosambi function because the Kosambi function incorporates crossover interference, the effect of which is absent in the mutants analyzed in this study (Figure 6D and table S2, and would have thus likely underestimated the genetic distances. The obtained recombination frequencies per interval and corresponding genomic data are shown in Table S4. Crossover interference patterns (Figure 6D) were analysed using MADpatterns^28^. Graphical representations were prepared with Graph Prism 6. Tests shown on figure 3A are one way Anova with Dunnet correction on the observed number of COs (Genotype transitions) per F2 plant in the genotyping data after error filtering. Male meiotic chromosome spreads have been performed as described previously^29^.

## Acknowledgments

We thank Ian Henderson for discussions and sharing data before publication. We are grateful to Christine Mézard, Mathilde Grelon and Eric Jenczewski for fruitful discussions and critical reading of the manuscript. We thank Gregory Copenhaver for providing the FTL lines. This work was funded by the European Research Council Grant ERC 2011 StG 281659 (MeioSight) and the Fondation Simone et Cino del DUCA/Institut de France. AHL was supported by the International Outgoing Fellowships PIOF-GA-2013-628128, POLYMEIO.

---

Table S1. References for data in Figure 1.

Table S2. Raw data of FTL tetrad analysis

Table S3. Raw genotyping data

Table S4. Genomic and recombination data

Table S5. Genotyping primers

